# Robin: An Advanced Tool for Comparative Loop Caller Result Analysis Leveraging Large Language Models

**DOI:** 10.1101/2025.04.05.646688

**Authors:** H. M. A. Mohit Chowdhury, Mattie Fuller, Oluwatosin Oluwadare

## Abstract

**Background:** There has been significant interest in genomics research, leading to the development of numerous new methods. One notable area of progress is in chromosome looping detection algorithms (also known as loop callers). However, despite these advancements, there is no available platform to analyze, compare, or benchmark current tools’ results on the go. Developing such a platform is crucial to accelerate research and ensure the reliability and effectiveness of new methods in the field.

**Results:** Hence, in this work, we propose Robin, an advanced ready-to-go platform for comparative loop caller result analysis leveraging Large Language Models (LLMs). Robin is a web server designed to analyze loop caller results, offering a comprehensive range of analysis metrics such as recovery and overlap. It is integrated with HiGlass and incorporates LLMs to enable users to generate plots simply by providing instructions.

**Conclusions:** Overall, Robin is a robust and comprehensive loop caller result analysis and visualization tool. It is publicly accessible at http://hicrobin.online, with a comprehensive documentation available at http://documentation.hicrobin.online/.

## 1 Background

Chromosome forms a complex and tightly confined structure by containing the DNA within its tiny compartment [1]. Inside a chromatin, loops are formed to fit into a small space and support DNA structure. These loops form in response to interactions both within and between chromosomes. Numerous proteins, including histone protein, SMC3, CTCF, and others are present in these loop regions [2]. Histone protein gives chromatin a compact structure, H3K27ac functions as a packaging protein, and CTCF is a transcriptional activator that inhibits communication between the enhancer and promoter [2]. These proteins also provide structural and functional support for chromatin. Scientists have been interested in these loop regions because of their diversity and biological significance.

There are lots of loop caller tools available such as cLoops [3], Mustache [4], HiC-CUPS [5], etc. with diversified input parameters and accepting different types of input data formats such as Hi-C [6], ChIP-seq [7], etc. Researchers often benchmark their tools by considering various use cases and employing different types of metrics. This process typically involves writing extensive boilerplate code, and there is currently no platform available to evaluate the performance of each tool comprehensively. To address this gap, Chowdhury et al. (2024) published the first comparative study of loop callers [8]. In this study, they analyzed and categorized twenty-two different types of loop callers, and benchmarked eleven of these tools side-by-side. The authors also introduced a novel Recovery Efficiency Metric (*REM*) score to demonstrate the tools’ efficiency in recovering specific proteins. This metric evaluates the performance of each loop caller by focusing on the recovery rate and normalizing it by the number of loops. This ensures fair, independent assessments of loop calling algorithms and prevents bias from tools that detect an excessive number of loops. [8].

Despite the valuable insights from this study [9], the analysis remains static and customized to the tools considered, leaving current and future tools analysis unsupported. Given the rapid growth in this field, there is a pressing need for a dynamic, on-the-go analysis tool that provides a comparative platform for ongoing research and supports the future development of loop calling methods. To address this need, we developed Robin, an advanced ready-to-go platform for comparative loop caller result analysis and visualization leveraging Large Language Model (LLM). Robin is a web server designed to analyze loop caller results, providing a comprehensive range of analysis metrics such as consistency, recovery, and regression analysis. Additionally, HiGlass [9] is integrated into its deployment. Robin leverages a LLM to facilitate the generation of user-specific plots with ease. Furthermore, users can download high-resolution plots directly from the Robin web server for their detailed analysis.

## 2 Implementation

Robin has five main features-Overlap, Regression Analysis, Recovery Visualization, HiGlass integration, and an AI data visualization assistant that leverages LLMs to generate new graphs at runtime. Overlap analysis shows the overlap of chromatin loops between up to three files of the same resolution in the form of a Venn diagram. Regression analysis shows both categorical (if categories were defined by the user) and linear regression of average size of the reported loops by caller (in Kb and number of bins) vs resolution. In this work, protein recovery plots display recovery and REM analysis, along with graphs comparing results at high resolutions of 5Kb and 10Kb, and low resolutions of 100Kb and 250Kb. It is worth noting that users can customize and define their own high and low resolution ranges based on their specific analysis needs. Robin’s HiGlass integration automatically processes all uploaded result files and protein reference files from the user. Additionally, it employs OpenAI’s GPT model [10] as a backend to generate scripts, presenting users with Jupyter Notebook files containing resulting graphs and source code during runtime.

### 2.1 Core Architecture

Robin consists of a few parts: i. frontend, ii. web-api or backend, iii. database. All of these three components are bundled together in various docker images, and they are connected in the same network and communicate with each other using the docker-compose command. Frontend is one of the main parts of Robin as users directly interact with this part. We used React with React-Bootstrap and used React-Styleguidist to generate an isolated development environment for each component and build developer documents. In this way, Robin’s web server was developed to be modular and scalable with most components being developed in isolation thus making side effects or unintended state changes less likely. We used React for its diversified features and it is also maintainable. We used chart.js to render our different types of plots. The web API handles all interactions among frontend, database, and HiGlass such as submitting jobs, retrieving data, communicating with AI assistants, and updating job information, etc. The web API is data-driven and executes analysis scripts to generate job results, act in response to the database, and run Jupyter Notebook. We used Python, R, and bash scrips to calculate loop size, overlap and recovery analysis. We also wrote a worker that handles Higlass, and converts and prepares data using bed-tools and HiGlass Clodious to make them compatible with the HiGlass environment. We implemented HiGlass and the main Robin server in different docker containers and Robin has the capabilities to communicate internally with the HiGlass server.

### 2.2 LLM System

The LLM is implemented in Robin’s AI Assistant by utilizing OpenAI’s API to generate Python code with a user-provided API key. The generated code is stored in the browser’s memory, allowing users to review and verify the AI-generated code. While the AI-generated Python code serves as an assistant, users are encouraged to thoroughly review it for correctness, as human oversight remains essential for catching potential logic errors. Once the user finds the provided code satisfactory and submits it, the code is sent to the Robin API and executed in a container running a Flask API to ensure it runs without any exceptions. After validation, the code is written to a Jupyter Notebook file, which is compiled and served to the end-user as an embedded HTML within the results page. These notebook files are stored in association with the job on Robin’s server, allowing all results to be viewed at any time.

## 3 Results

Robin is a web server for comparative loop caller result analysis leveraging LLM model (Figure 1). Users can upload their loop caller results to Robin and export the analytical result plots in high resolution. It is integrated with HiGlass [9]. HiGlass is a popular server for visualizing signals. We have seamlessly integrated HiGlass for Robin users, streamlining the process and making it easy-to-use for variety of users. Below we describe an example result-set of the Robin for loop caller result analysis similar to the analysis performed and reported in the comparative study by Chowdhury et al. using Human Lymphoblastoid cells (GM12878) dataset [8].

**Fig. 1.**
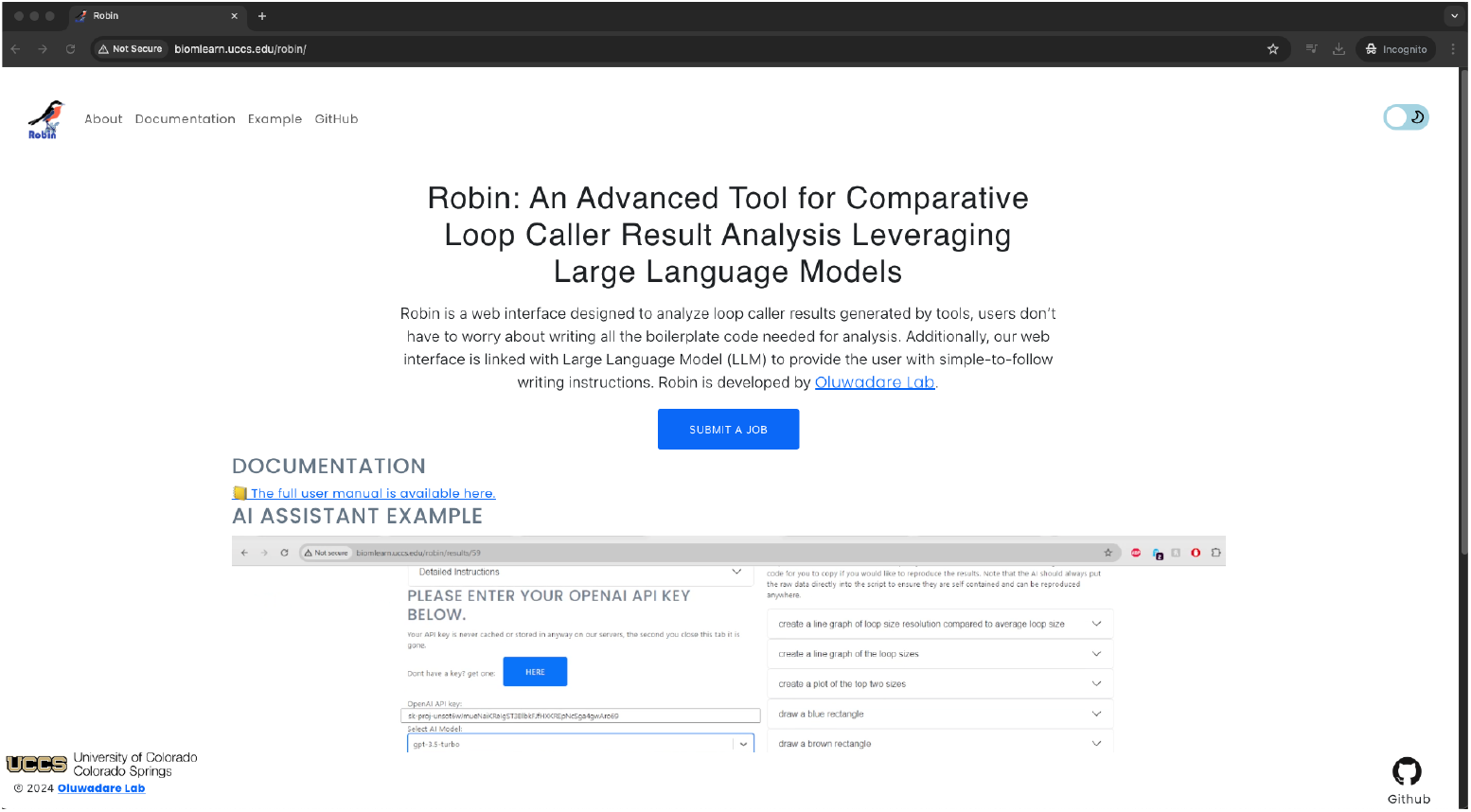
Robin web server landing page for loop caller result analysis. From this page users can navigate to different options such as submit a job, reading documentation etc.

### 3.1 Primary result

The first tab of results reports on the detected overlaps (Figure 2A). This overlap showcases the percentage of commonly identified loop regions, elucidating the consistency of loop callers across up to three tools in a single plot. Users can navigate through different resolutions. Following is the regression analysis plot tab, which includes plots of Average Bin Size (number of bins and size in Kb) versus Resolution (Figure 2B). These plots illustrate how the bin size changes with resolution and display the average number of bins at a specific resolution. This tab also contains regression plots (both overall and categorical) to demonstrate the consistency of the loop callers. Subsequently, the protein recovery result tab is presented in Figure 2C. Recognizing biological features as paramount compared to others, we offer various types of protein-specific (CTCF, H3K27ac and RNAPII) recovery rate plots (both line and bar plots) according to user specifications. In the protein recovery rate line plot, x-axis describes the number of loops predicted. Every loop predicts a fixed number of loops, our analysis segments them into every 1000 loops count interval in an increasing order. Following x-axis, y-axis presents the Recovery Rate at every 1000 loops count increasingly. If a user chooses a random point in the graph, it explains the recovery rate out of the first x-amount of loops predicted by a loop caller tool and the edge point of every tool indicates the total recovery rate out of total loops predicted by a specific loop caller. Additionally, for logical comparison, we incorporate the REM plot, showcasing the efficiency of each tool’s recovery rate over the detected loop count. To illustrate consistency between two resolutions, we include a consistency plot comparing high (5Kb, 10Kb) and low (100Kb, 250Kb) resolutions by averaging the REM score. We included three protein (CTCF, H3K27ac and RNAPII) analysis results in our example analysis (Figure 2C) to show how this feature works, however, our platform allows users to upload as many proteins as needed for further analysis and every protein specific tab will be shown as exemplified in our example analysis. HiGlass provides the ChIP signal plots of every individual tool (Figure 2D). It shows the real positional alignment of protein-specific positions with the original protein reference signals. It helps to visualize and validate recovery plots. Moreover, we included Data tab to show users uploaded data for further uses with download option (Figure 2F).

**Fig. 2.**
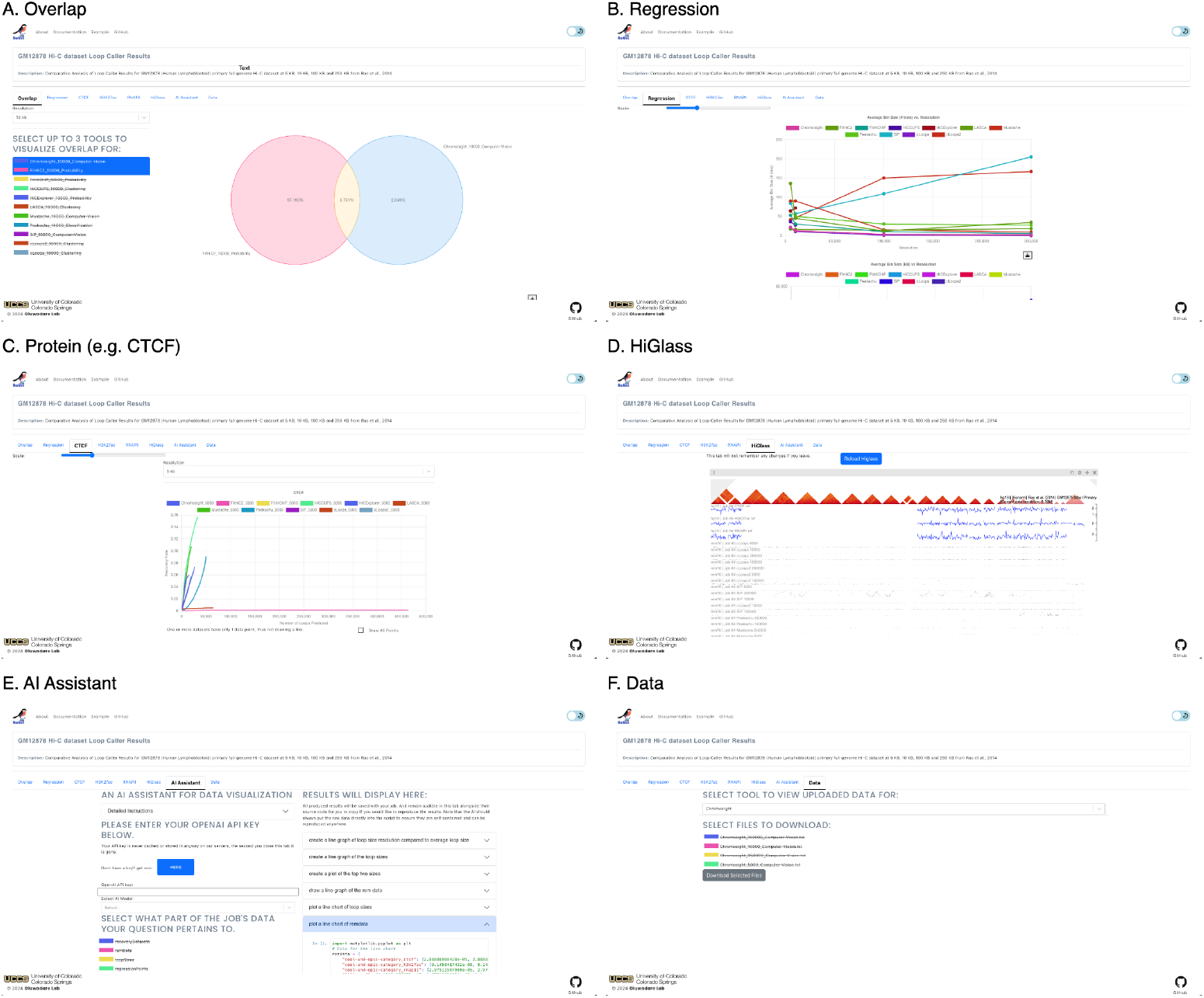
Analysis results using different loop-caller results at high (5Kb, 10Kb) and low (100Kb, 250Kb) resolutions. (A) Robin’s overlap tab contai ning the Venn diagram of tools’ result, (B) Robin’s regression tab containing Average Bin Size vs Resolution and Regression plots, (C) Robin’s protein specific tabs named according to uploaded protein reference file, and contains recovery rate vs. number of loops and REM plots, (D) Integrated HiGlass tab to visualize ChIP signals along with loops results, (E) AI Assistant tab to generate custom plots not available in Robin by default, using the uploaded dataset based on the user’s simple commands, (F) User uploaded data for further download.

### 3.2 Hybrid Analysis: User-Supervised LLM Generated Code

Leveraging the LLM (OpenAI GPT) allows the user more freedom and options for quick data analysis and visualization. The AI Assistant included in the results tab (Figure 2E), allows the user to prompt the AI system to generate new graphs to visualize data analysis that is not shown in Robin. This process begins with instructing the AI to generate code snippets for performing a desired analysis based on the data uploaded by the user. Once the code is generated, the user reviews and supervises the code to ensure logical correctness, making any necessary updates before submitting it to be executed within Robin. All AI-generated code is fully self-contained, including the data it references, which allows these scripts to be easily edited on the Robin platform and downloaded for use elsewhere without requiring additional data files.

As previously mentioned, user supervision is a crucial component of this process. While LLMs can generate code effectively, it is important to recognize that they may occasionally produce results containing errors or missing logical components. Therefore, while the AI Assistant is a valuable tool for rapidly prototyping data visualizations, we strongly advise users to thoroughly review and supervise the generated code, using it for exploratory analysis. With appropriate user oversight, this tool can greatly facilitate quick and exploratory data analysis and visualization of user-uploaded loop caller results, extending the capabilities beyond what Robin natively provides.

For example, in our case study, we instructed the AI Assistant to perform the following analyses: create a line graph comparing loop size resolution to average loop size, generate a line graph of the loop sizes, and produce a plot of the top two loop sizes, among others. Once the code for these instructions was generated, we utilized Robin’s built-in editor to review and refine the code as needed. After completing this review, we submitted the code within Robin to produce the requested figures (Figure 2E). The backend mechanics of the LLM system are detailed in Section 2.2.

We have provided comprehensive documentation and an example job on the Robin website to assist users in utilizing the web server.

## 4 Conclusions

Robin represents a significant advancement in chromatin loop caller analysis, offering a readily accessible platform complete with analysis metrics and a user-friendly web server. Users can leverage Robin-generated analysis plots, thereby preserving plot quality and accessing standard and scalable benchmarking metrics as extended from Chowdhury et al. (2024) [8]. Integration of HiGlass visualization tools within Robin facilitates the comparison of ChIP signals for assessing consistency among tools. Additionally, the incorporation of LLM integration expands the scope of Robin’s utility for users. Through the AI assistant, users can effortlessly generate plots not provided by Robin with a single sentence, utilizing the same dataset. Furthermore, Robin equips users with interactive plots and the option to download high-resolution plots for further analysis. Over all, Robin offers an intuitive and robust analysis platform for loop caller algorithms, thereby streamlining the research process.

## 5 Availability and requirements

**Project name**: Robin

**Project home page**: http://hicrobin.online

**Operating system(s)**: Platform independent

**Programming language**: Python, R, JavaScript, HTML, CSS

**Other requirements**: Docker, Web Browser

**License**: GNU GPL v3

**Any restrictions to use by non-academics**: None

## 6 Declarations

### 6.1 Ethics approval and consent to participate

Not applicable.

### 6.2 Consent for publication

Not applicable.

### 6.3 Availability of data and materials

We used Human Lymphoblastoid (GSE63525 GM12878) cell to generate the example usecases included in Robin’s example page. All our backend and documentation code are available at https://github.com/OluwadareLab/Robin and https://github.com/OluwadareLab/Robin documentation respectively.

### 6.6 Authors’ contributions

HMAMC wrote the backend analysis codes. MF developed the webserver, and OO conceived and supervised this project. All authors wrote and reviewed the manuscript.

## 6.7 Acknowledgements

The authors acknowledge Drew Houchens from the Bioinformatics Lab for his contribution towards the logo design. The authors acknowledge Abhishek Pandeya, Samuel Olowofila, Lin Lin, Mohan Baleathiguppe Chandrashekar, Jahanara Mohamed from Bioinformatics Lab for their contribution towards testing and feedback to improve the user experience of Robin.

